# Accuracy of serological tests for diagnosis of chronic pulmonary aspergillosis: a systematic review and meta-analysis

**DOI:** 10.1101/763094

**Authors:** Cláudia Elizabeth Volpe Chaves, Sandra Maria do Valle Leone de Oliveira, James Venturini, Antonio Jose Grande, Tatiane Fernanda Sylvestre, Rinaldo Poncio Mendes, Anamaria Mello Miranda Paniago

## Abstract

Chronic pulmonary aspergillosis (CPA) is a disease that benefits from cavities as after-effects of tuberculosis, presenting a high mortality rate. Serological tests like double agar gel immunodiffusion test (DID) or the counterimmunoelectrophoresis (CIE) test have been routinely used for CPA diagnosis in the absence of positive cultures; however, they have been replaced by enzyme-linked immunoassay (ELISA), with a variety of methods.

This systematic review aims to compare the accuracy of the ELISA test with the reference test (DID and/or CIE) in CPA diagnosis. It was conducted according to the Preferred Reporting Items for Systematic Reviews and Meta-Analyzes (PRISMA).

The study was registered in PROSPERO under the registration number CRD42016046057. We searched the electronic databases MEDLINE (PubMed), EMBASE (Elsevier), LILACS (VHL), Cochrane library, and ISI Web of Science. Gray literature was researched in Google Scholars and conference abstracts. We included articles with patients or serum samples from CPA patients who underwent two serological tests: ELISA (index test) and IDD and/or CIE (reference test), using the accuracy of the tests as a result. Original articles were considered without a restriction of date or language. The pooled sensitivity, specificity, and summary receiver operating characteristic curves were estimated.

We included 13 studies in the review, but only four studies were included in the meta-analysis. The pooled sensitivities and specificities were 0.93 and 0.97 for the ELISA test. For the reference test (DID and/or CIE), these values were 0.64 and 0.99. Analyses of summary receiver operating characteristic curves yielded 0.99 for ELISA and 0.99 for the reference test (DID and/or CIE). Our meta-analysis suggests that the diagnostic accuracy of ELISA is greater than that of the reference tests (DID and/or CIE) in early detection of CPA.

## Introduction

Chronic pulmonary aspergillosis (CPA) is a slow and progressive lung disease caused by *Aspergillus spp.* that develops in preexisting cavities of patients with chronic respiratory diseases, and pulmonary tuberculosis is its main predisposing factor, with a global prevalence estimated at 1.2 million cases [1].I Its prognosis is poor, with 38-85% mortality in five years [1, 2].

CPA presents five clinical forms: 1. aspergillus nodule, 2. pulmonary simple aspergilloma, 3. chronic cavitary pulmonary aspergillosis (CCPA), also called complex aspergilloma, 4. chronic fibrosing pulmonary aspergillosis (CFPA), and 5. subacute invasive pulmonary aspergillosis (SAIA) [3]. Aspergilloma is present in only one-third of patients with CPA [1, 4].

The diagnosis of CPA is based on suggestive images, preferably tomographic images (CT scan), on evidence of microbiological infection by *Aspergillus* or on the presence of an immune response to this agent, maintained for at least 3 months [3, 5].

Serologic tests are indispensable for the diagnosis in the absence of positive cultures and are considered the best noninvasive tests to diagnose this entity [6, 7]. These tests may be over 90% positive with precipitins or in the detection of *Aspergillus* IgG [2, 3].

In patients presenting *Aspergillus* in the respiratory tree, the detection of specific serum antibodies differentiates infection from colonization, with a positive predictive value of 100% for identification of infection [8]. Initially, antibodies against *Aspergillus fumigatus* were determined by detection of precipitins using the double agar gel immunodiffusion test (DID) or the counterimmunoelectrophoresis technique (CIE) [4,9,10] with a sensitivity of 89.3% [6] and a specificity of 100% [11].

These methods (DID and CIE) consume a lot of time, intense work, require relatively large extracts of *A. fumigatus* and patient serum, and provide only semiquantitative results [7].

The *Aspergillus* IgG antibody test is strongly recommended by the Infectious Diseases Society of America IDSA [12]. In practice, precipitation techniques have already been replaced by the *Aspergillus* IgG antibody detection test by enzyme-linked immunoassay (ELISA) [13]. This is considered the fastest and most sensitive test [14], producing quantitative results with lower extracts of *A. fumigatus* and patient serum by test, besides it is easily automated [7].

Despite its importance, serology for the detection of *Aspergillus* IgG by ELISA still does not reach a definitive conclusion on diagnostic performance for CPA, as significant differences in sensitivity, specificity and coefficient of variation need to be explored with cohorts of well-characterized patients [3].

Historically, IgG ELISA assays used in-house antigens, with different antigenic preparations and concentrations, which makes the comparison of test performance very difficult [7]. Currently, we have commercial tests such as ELISA plates for *Aspergillus*-specific IgG antibodies produced by Serion (Germany), IBL (Germany / USA), Dynamiker / Bio-Enoche (China), Bio-Rad (France), Bordier (Switzerland) and Omega / Genesis (UK), as well as specific *Aspergillu*s IgG automated systems such as Immunolite-Siemens (Germany) and ImmunoCAP (Thermo Fisher Scientific / Phadia), which are fluoroenzyme immunoassay variants of ELISA. The main limitation of these tests is the detection of antibodies only against *A. fumigatus* and as they account for only 40% of the isolates, diagnosis of CPA caused by non-*fumigatus* strains is still a challenge [2].

Considering the variety of methods for detection of antibodies to *Aspergillus*, the use of precipitation tests due to their low cost and the absence of more precise options for serological diagnosis of CPA, the present study review on serological diagnosis of chronic pulmonary aspergillosis, comparing the performance of the precipitation tests with the enzyme-linked immunoassay tests.

## Materials and Methods

We conducted a systematic review of the literature in accordance with the recommendations of the Preferred Reporting Items for Systematic Reviews and Meta-Analyzes (PRISMA) [15] and STARD 2015 [16]. A protocol for systematic review was developed and registered in the International Prospective Register of Systematic Reviews - CRD42016046057. We used the Cochrane recommendations to report systematic reviews and meta-analyses of studies on diagnostic accuracy [17].

### Eligibility criteria

We considered as inclusion criteria articles with population or serum samples from patients diagnosed with aspergilloma or chronic pulmonary aspergillosis that were submitted to the ELISA immunoenzymatic test (ELISA test) and to the double immunodiffusion gel agar and/or counterimmunoelectrophoresis test (DID and/or CIE). The accuracy of the tests was defined as primary outcome. Original studies were included without restriction of language, geographical location or date of publication. We excluded studies with children or animals and/or in *vitro*. We were unable to find an article in Japanese, which was selected for full article reading because it was not available in the international library commuting service.

### Information sources and search strategies

The studies were searched in the following databases: MEDLINE (through PubMed), EMBASE (through Elsevier), LILACS (through VHL), Cochrane library and ISI Web of Science. Gray literature was researched in Google Scholars and congress abstracts. We submitted the search strategy performed until June 2019.

We used the following search strategy for Medline and adapted it for the other databases: pulmonary aspergillosis AND serologic test (and its synonyms). 1. ((“Pulmonary Aspergillosis” [Mesh] or Aspergillosis, Pulmonary or Pulmonary Aspergillosis or Lung Aspergillosis or Aspergillosis, Lung or Aspergillosis, Lung or Bronchopulmonary Aspergillosis or Aspergillosis, Bronchopulmonary or Bronchopulmonary Aspergillosis or Aspergillosis, Bronchopulmonary or Aspergillose, Bronchopulmonary or Bronchopulmonary Aspergillose) AND (“Serologic Tests” [Mesh] or Serological Tests or Serological Tests or Serological Tests, Serological or Tests, Serologic or Serologic Tests or Serologic Tests or Serodiagnoses).

### Study selection and data extraction

Titles were imported from EndNote Online and duplicate studies were removed. The remaining titles were independently reviewed by two authors (TFS and SMVLO), who selected the article abstracts, as well as defined the complete texts for evaluation. The divergences were resolved by a third expert reviewer (RPM). Two other authors (CEVC and JV) performed independent evaluations of the complete articles and judged the methodological quality of the included studies using the Quality Assessment of Diagnostic Accuracy Studies (QUADAS-2) tool [18]. The divergences were resolved by consensus among the researchers.

Two reviewers (CEVC, JV) independently extracted the following data from each included study:

- Study characteristics: author, year of publication, country, design, and sample size.
- Population characteristics: according to the inclusion criteria
- Description of the index test and cut-off points;
- Description of the reference standard and cut-off points;
- QUADAS-2 items;
- Accuracy results obtained in each study to construct a diagnostic contingency (two-by-two table);

### Assessment of methodological quality

For this review, we used the QUADAS-2 tool to assess the methodological quality of studies [18]. QUADAS-2 consists of four key domains: patient selection, index test, reference standard, and flow and timing. We assessed all domains for the potential of risk of bias (ROB) and the first three domains for concerns regarding applicability. Risk of bias is judged as “low”, “high”, or “unclear”. Two review authors independently completed QUADAS-2 and resolved disagreements through discussion.

### Statistical analysis and data synthesis

We used data reported in the true positive (TP), false positive (FP), true negative (TN) and false negative (FN) format to calculate sensitivity and specificity estimates and 95% confidence intervals (CIs) for individual studies. Summary positive (LR+) and negative (LR-) likelihood ratios and summary diagnostic odds ratio (DOR) were obtained from the bivariate analysis. We used the clinical interpretation of likelihood ratios [19] as follows: conclusive evidence (LR+>10 and LR-<0.1), strong diagnostic evidence (LR+ >5 to 10 and LR- 0.1 to <0.2), weak diagnostic evidence (LR+ >2 to 5 and LR- 0.2 to <0.5) and negligible evidence (LR+ 1 to 2 and LR- 0.5 to 1).

In studies where it was possible to calculate sensitivity and specificity for the ELISA test and DID and/or CIE, we calculated accuracy test and Youden’s J statistic. The Youden’s index values range from zero to one inclusive, with the expectancy that the test will show a greater proportion of positive results for the diseased group than for the control [20].

Studies were submitted to meta-analysis when three conditions were required: 1. sample size greater than 20; 2. sensitivity and specificity were available for the index and the reference tests; 3. healthy controls were included in the analysis. We presented individual studies and pooled results graphically by plotting the estimates of sensitivity and specificity (and their 95% CIs), heterogeneity and receiver operating characteristic (ROC) space using Stata software. For the subgroup analysis we presented individual studies and pooled results in forest plots using Meta-DiSc software.

### Investigations of heterogeneity

We investigated heterogeneity by subgroup analyses. We addressed the main source of heterogeneity: in-house and commercial ELISA tests. In-house tests have presented many technical differences. We considered an I2 value close to 0% as having no heterogeneity between studies, close to 25% with low heterogeneity, close to 50% with moderate heterogeneity and close to 75% with high heterogeneity between studies [21].

## Results

### Study inclusion

A total of 2096 articles were identified in five databases, of which 2010 were searched through a database and 63 articles were identified from other sources (manual search). After the removal of duplicates, we remain with 1797 articles. After title / abstract exclusion, only 20 articles were submitted to a full text read and 13 of them were included for the systematic review; only four studies were included for the meta-analysis (see Fig 1).

**Fig 1.**
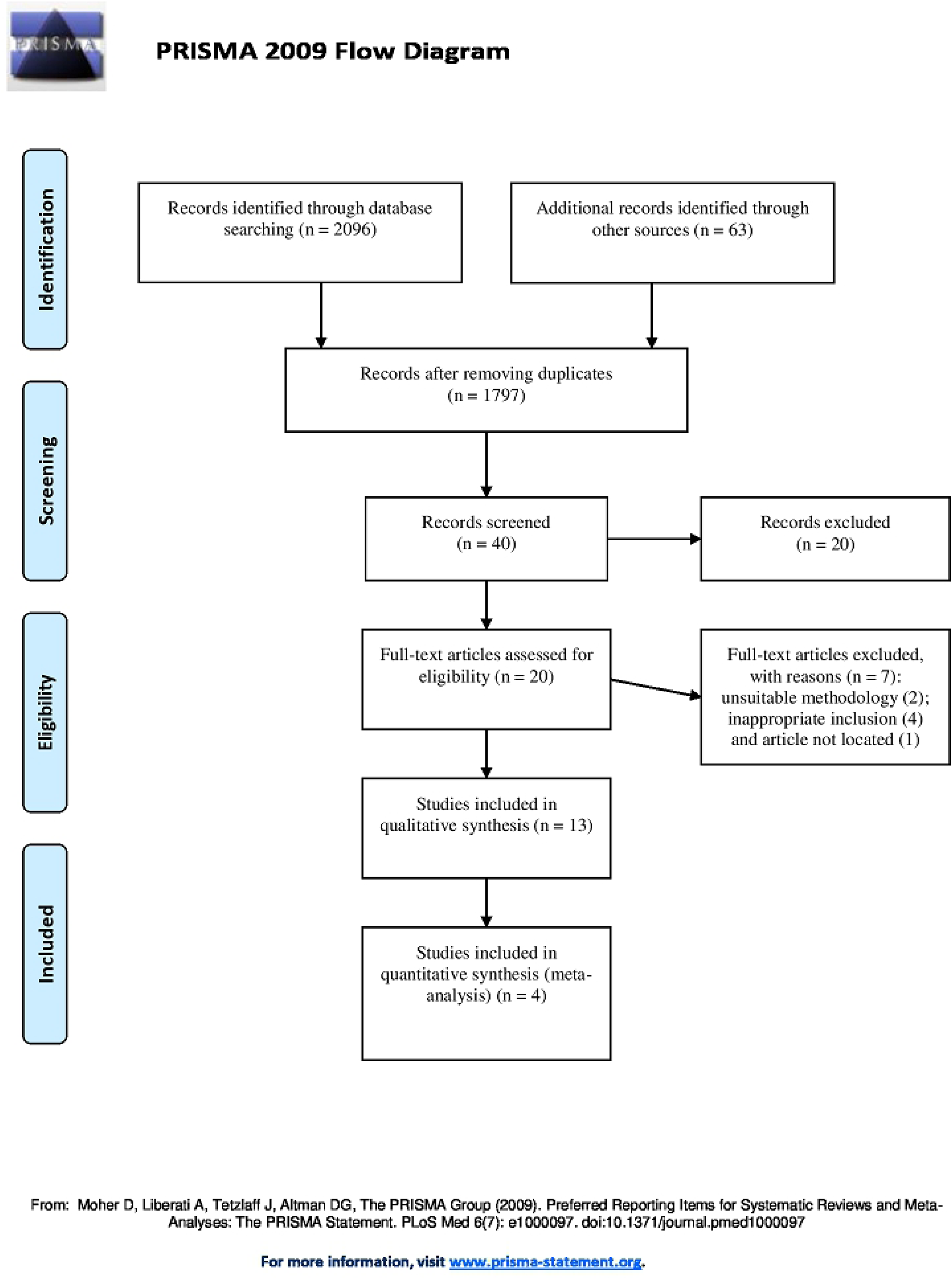
Study flow diagram.

### Characteristics of the studies

The characteristics of the included studies are presented in S1Table. The earliest study was published in 1983 [22] and the five most current articles were published in 2015 [23], 2016 [24, 25, 26] and 2018 [27]. Nine studies took place in five countries: Japan [25, 28], Brazil [23], United Kingdom [24, 29], France [26, 30, 31] and India [27], but in 4 articles, the study countries were not reported [22, 32, 33, 34].

Nine articles presented DID as the reference test [22, 23, 25, 27, 28, 30, 32, 33, 34]; an article presented two reference tests, DID and CIE [34] and four studies presented only CIE as the reference test [24, 26, 29, 31].

When we performed data extraction, some important differences were observed and deserve to be highlighted. Seven articles performed in-house ELISA tests [22, 23, 28, 30, 32, 33, 34] and six articles described their studies with commercial tests [24, 25, 26, 27, 29, 31]. Different *Aspergillus* antigens and cut-off points were used in the in-house ELISA tests; the articles that used commercial tests also used several types of antigens and cutoff points included by authors beyond those established by the manufacturer and are described in S1 Table.

In one article, we were unable to identify the number of patients evaluated with CPA, nor was it possible to extract data from the 2×2 table for DID and ELISA [28]; in two articles it was not possible to recover the DID data [25, 30]; in another article, data were not obtained from CIE [31] and in another [32], it was not possible to extract data for ELISA. In one study [33], 20 sera from 13 patients were used and it was not possible to extract the accurate data per patient, besides data from the control group was not presented for the ELISA test; in two articles, the tests were not submitted to a control group [26, 29]; furthermore, in one article, the control group was performed on patients with any presence of DID precipitation line and it was not considered by us as a control group [25].

During the extraction of ELISA antigen concentration data, five studies with in-house tests presented concentrations varying from 0.1 mcg to 250 mcg per well [22, 23, 30, 33, 34]; in two articles these concentrations were not reported [28, 32].

In the in-house tests, we still find other differences, such as ELISA secondary antibody dilution, with concentrations ranging from 1: 100 to 1: 300 when they were described [22, 23, 33, 34]; in three articles these dilutions were not reported [28, 30, 32]. When we evaluated the cut-off for ELISA, several descriptions were found with titers ranging from 1: 100 to 1: 800; we also found values in OD (optic density), au / mL, in percentage and in absorbance, and there was no comparable value in in-house tests [22, 23, 33, 34]; in three articles, the cut-off was not described [28, 30, 32]. For the ELISA substrate, TMB (3,3′,5,5′-Tetramethylbenzidine) was found in two articles [22, 23], also pNPP (Alkaline Phosphatase Yellow) [33, 34] and OPD (o-Phenylenediamine) [30]; in two articles the substrate was not reported [28, 32].

When extracting antigen concentration data from *Aspergillus fumigatus* in the studies for DID or CIE, we found variations between 5mg / mL and 100mg / mL[22, 29, 32, 33, 34]; we found values expressed in microliters in the following studies: 2 μL[31], 10 μL[26] and 20 μL[24]; and in one article different concentrations were used for somatic antigen [20 mg / mL] and antigen filtration [2mg / mL) [29]. The DID concentrations were not described in three articles [23, 28, 30].

The studies with commercial ELISA tests used the following tests: ImmunoCap [29, 24, 27, 25], Platelia [29], Immulite [24], Serion [24, 31], Dynamiker [24], Genesis [24], Bio-Rad [31, 26], and Bordier [26]. These tests presented different cut-off points and the one with the best performance is described in S1 Table.

All methodological differences can be observed in S1 table.

### Risk assessment of bias

We illustrated the methodological quality of the included 13 studies using the QUADAS-2 tool (Figs 2 and 3). All studies had unclear or high risk of bias in at least one domain. Almost all studies [22, 23, 24, 25, 27, 28, 29, 30, 31, 32, 33, 34] demonstrated high-risk patient selection bias, except one that was unclear (26), resulting mainly from not using consecutive or randomized patient samples and not avoiding a case-control study. In seven studies [22, 28, 30, 31, 32, 33, 34], there is not a clear definition of exclusion criteria.

**Fig 2.**
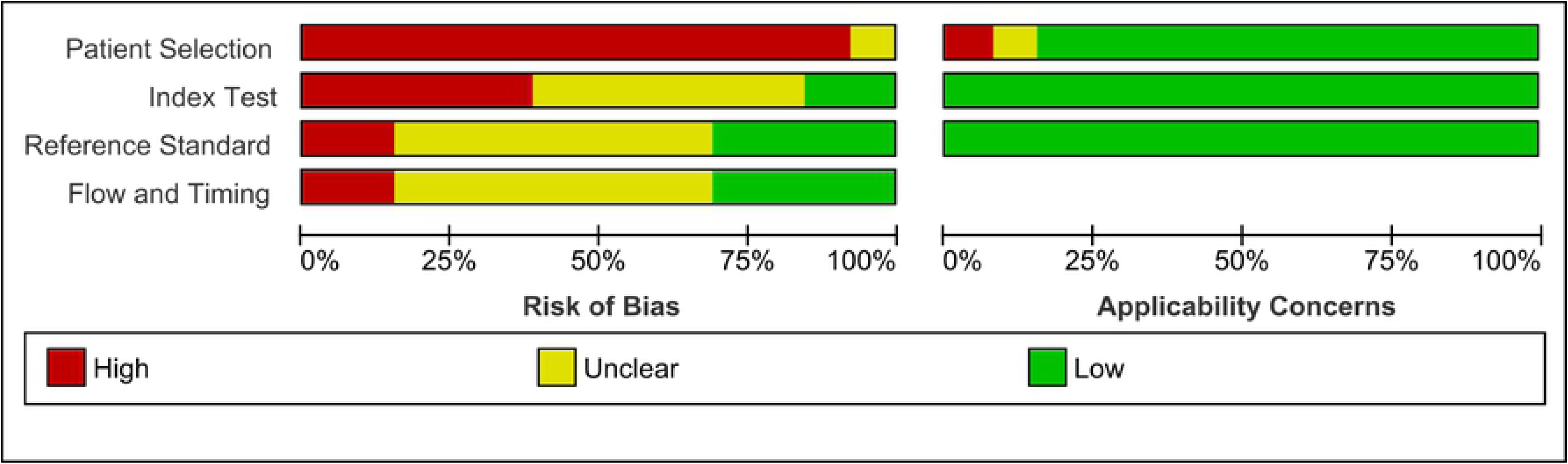
Proportion graph of studies assessed as having low, high or unclear risk of bias and/or applicability concerns.

**Fig 3.**
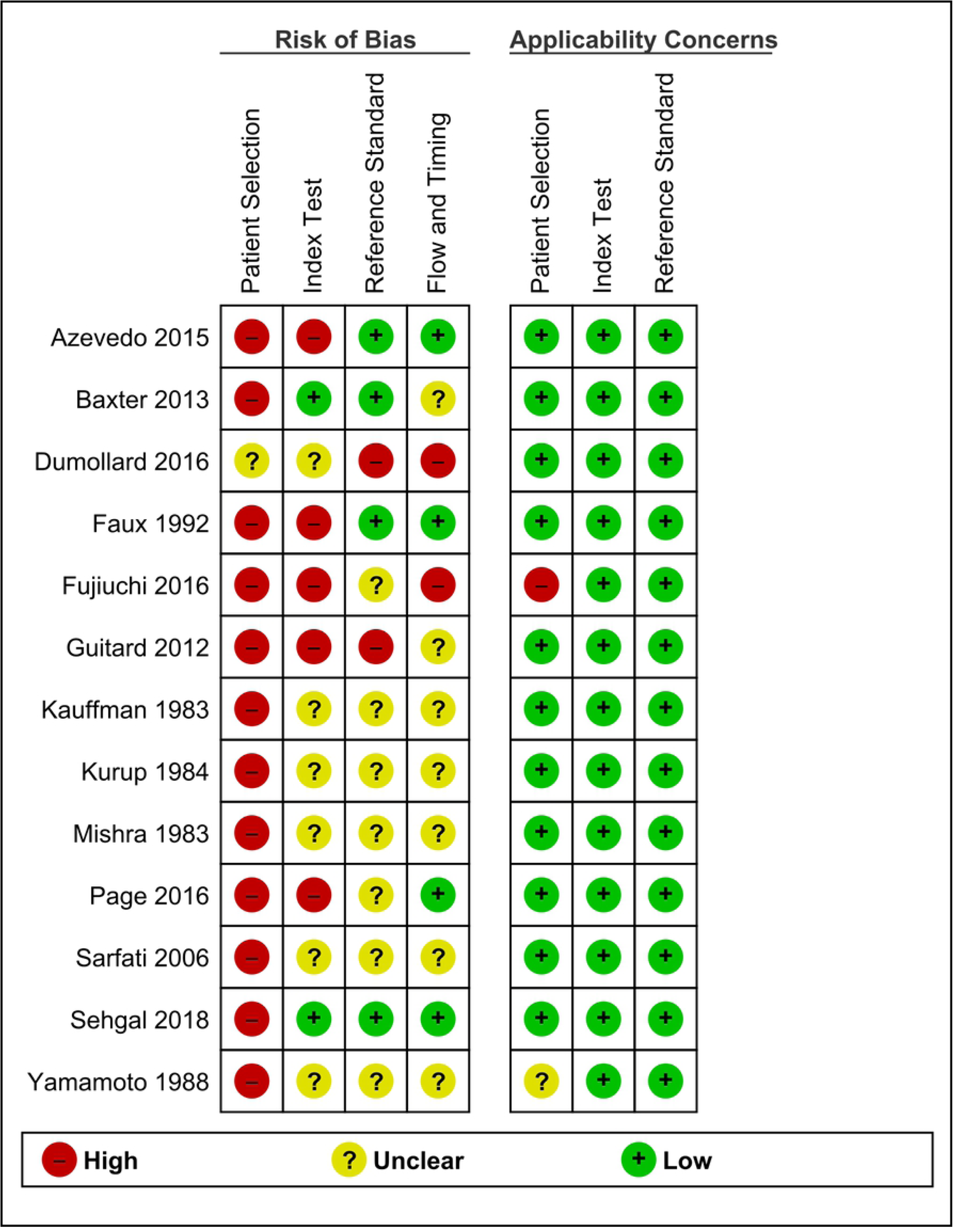
Risk of bias and applicability concerns graph: review of the authors’ judgments about each domain presented as percentages across included studies.

In the index test, eleven studies [22, 23, 24, 25, 26, 28, 30, 31, 32, 33, 34] presented an unclear or high risk of bias; mainly because the index test was interpreted with prior knowledge of the standard test. Eleven studies had a low risk of bias in the previous cut-off determination [22, 23, 24, 25, 26, 27, 28, 29, 31, 33, 34].

In the reference test, all studies had a low risk of correctly classifying the target condition; bias risk assessment was uncertain or high risk in 9 studies [22, 24, 25, 26, 28, 30, 31, 33, 34] for not making it clear whether the standard test was interpreted without the knowledge of the index test or if they already had prior knowledge.

Regarding flow and time, bias risk assessment was uncertain in eight studies [22, 26, 28, 29, 30, 31, 33, 34] for not clearly describing whether there was an appropriate interval between conducting the index test and the reference test; in one study [25] the evaluation was high risk. In eleven studies, all patients were submitted to a reference test, it was included in the analysis [22, 23, 24, 25, 27, 29, 30, 31, 32, 33, 34] and they had low risk; in one study, not all patients were submitted to a test reference [26] and in one study [28] this was uncertain.

Regarding applicability, almost all the articles presented low concern, because they did not fail to correspond to the critical question of this study.

### Diagnostic accuracy

We present the Table 1 with all the articles included in this systematic review, with a description of the index and reference tests, a number of patients and healthy controls, and a presentation of the values of sensibility, specificity, accuracy test, likelihood positive value, likelihood negative value and Youden’s statistic.

**Table 1.**
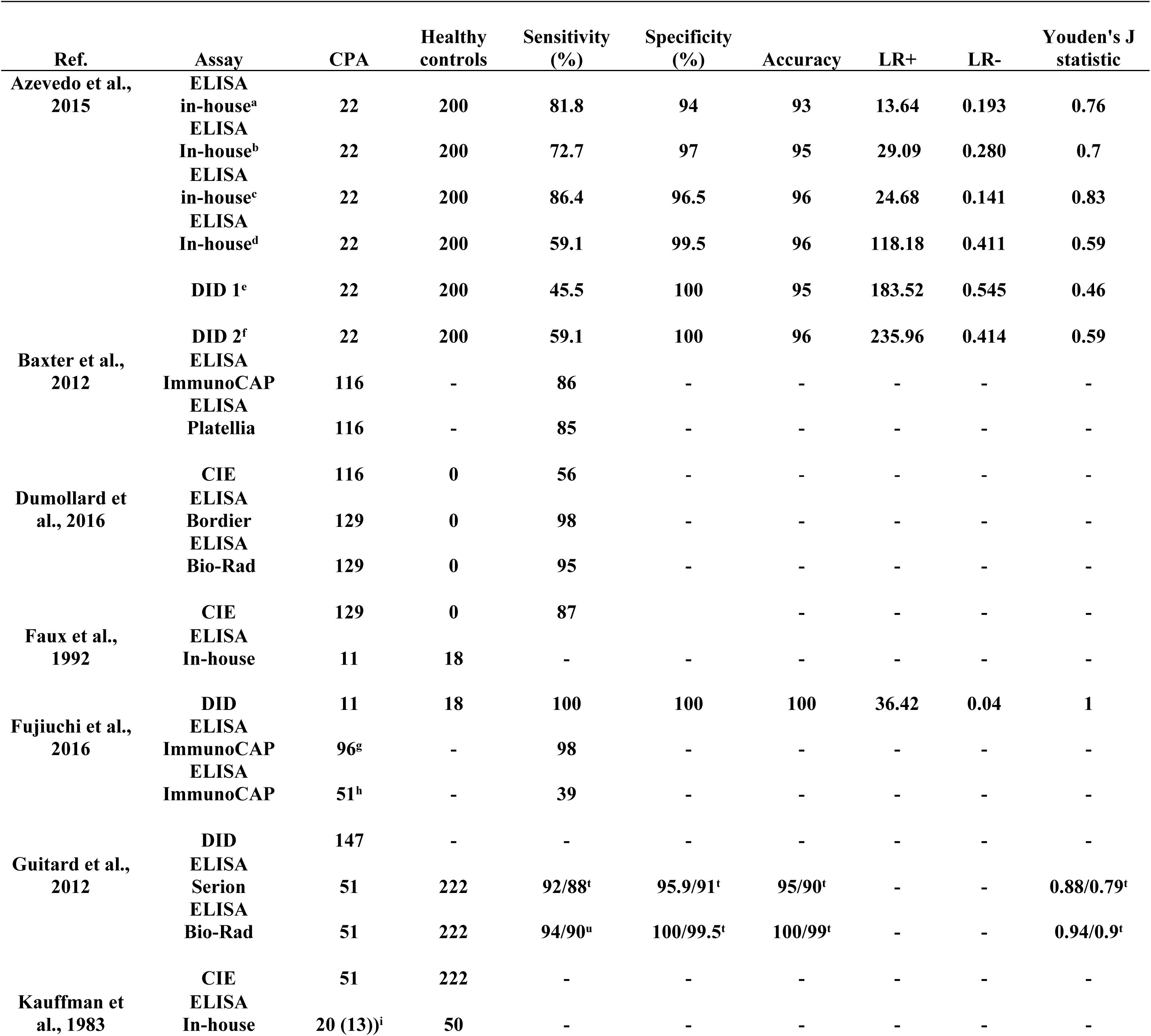
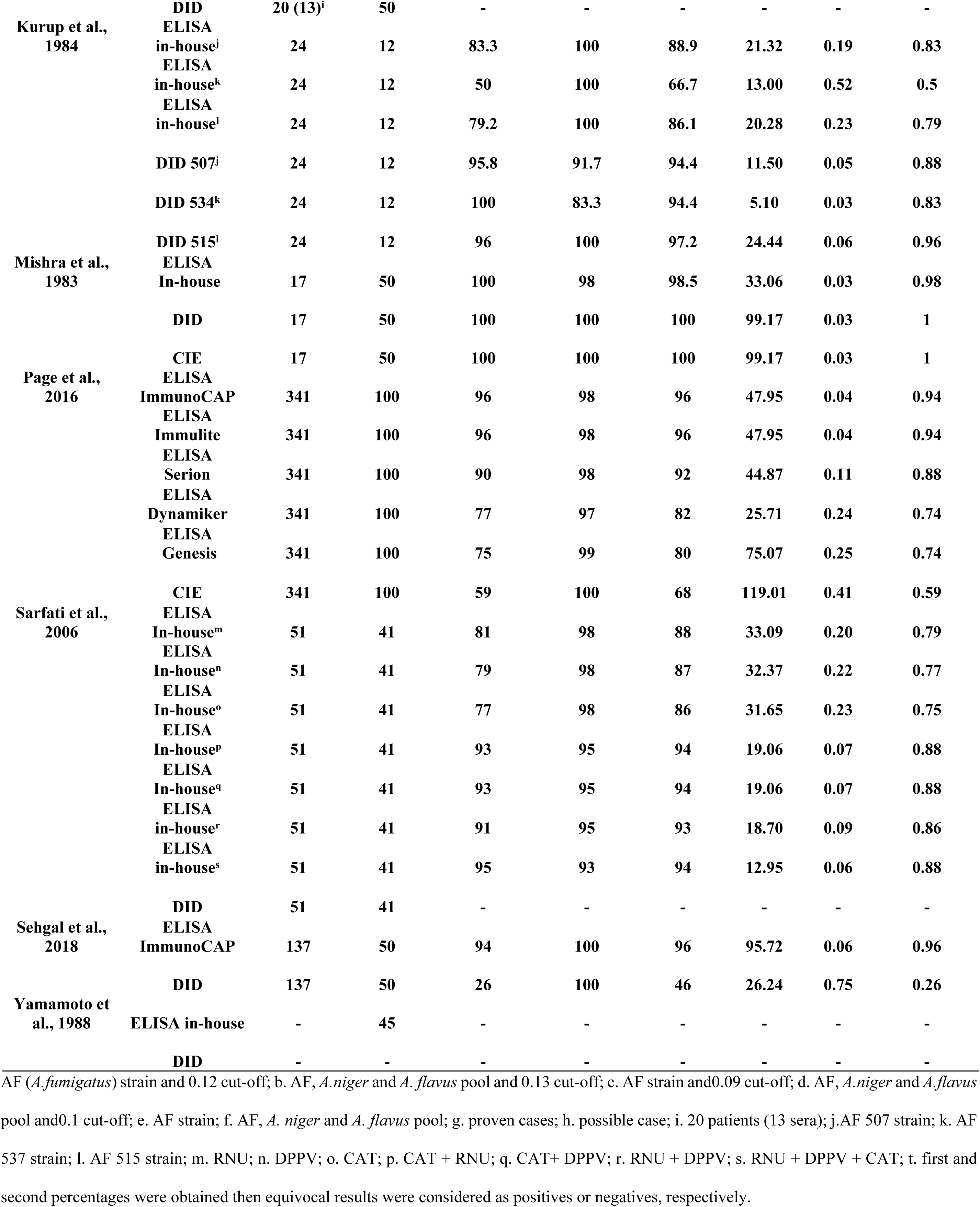
Performance of ELISA test and reference tests in studies included in systematic review.

The Youden index ranged from 0.50 to 0.98 for the ELISA test and from 0.26 to 1 for the reference test (DID and/or CIE) for the individual studies. Three studies presented a good performance above 0.90 Youden index for the reference test [22, 32, 34]. The other studies presented a performance below 0.90. The Youden indicates the trade-off between sensitivity and specificity.

### Quantitative synthesis - meta-analysis

In individual studies included in the meta-analysis, ELISA test sensitivity ranged from 0.83 (95% CI 0.63 to 0.95) [22] to 0.96 (95% CI 0.93 to 0.98) [24] and specificity ranged from 0.92 (95% CI 0.64 to 1.00) [22] to 0.98 (95% CI 0.93 to 1.00) [24]. The pooled sensitivity and specificity for the ELISA test, based on four data studies [22, 23, 24, 27], were 0.93 (95% CI 0.87 to 0.96) and 0.97 (95% CI 0.94 to 0.98), respectively. Pooled LR+ and LR- were 31.40 (95% CI 16.40 to 60.10) and 0.07 (95% CI 0.04 to 0.14), respectively. Pooled DOR were 440.00 (95% CI 156.00 to 1241.00). We interpreted the pooled LR+/LR- from the ELISA test as conclusive evidence, but we have not interpreted the reference test (DID and/or CIE) in the same way, because LR- was included as weak diagnostic evidence.

In the DID and/or CIE tests analyses, the sensitivity and specificity in individual studies ranged from 0.26 (95% CI 0.18 to 0.34) [27] to 0.96 (95% CI 0.79 to 1.00) [22] and 0.92 (95% CI 0.64 to 1.00) [22] to 1.00 (95% CI 0.97 to 1.00) [23], respectively. The pooled sensitivity and specificity for DID and/or CIE tests were 0.64 (95% CI 0.29 to 0.89) and 0.99 (95% CI 0.96 to 1.00). Pooled LR+/LR- were 53.00 (95% CI 19.20 to 146.40) and 0.36 (95% CI 0.14 to 0.92). Pooled DOR were 146.00 (95% CI 40.00 to 532.00).

The forest plots in Figs 4 and 5 show the sensitivity, specificity ranges and heterogeneity for the ELISA test and reference test (DID and/or CIE) in detecting chronic pulmonary aspergillosis across the included studies.

**Fig 4.**
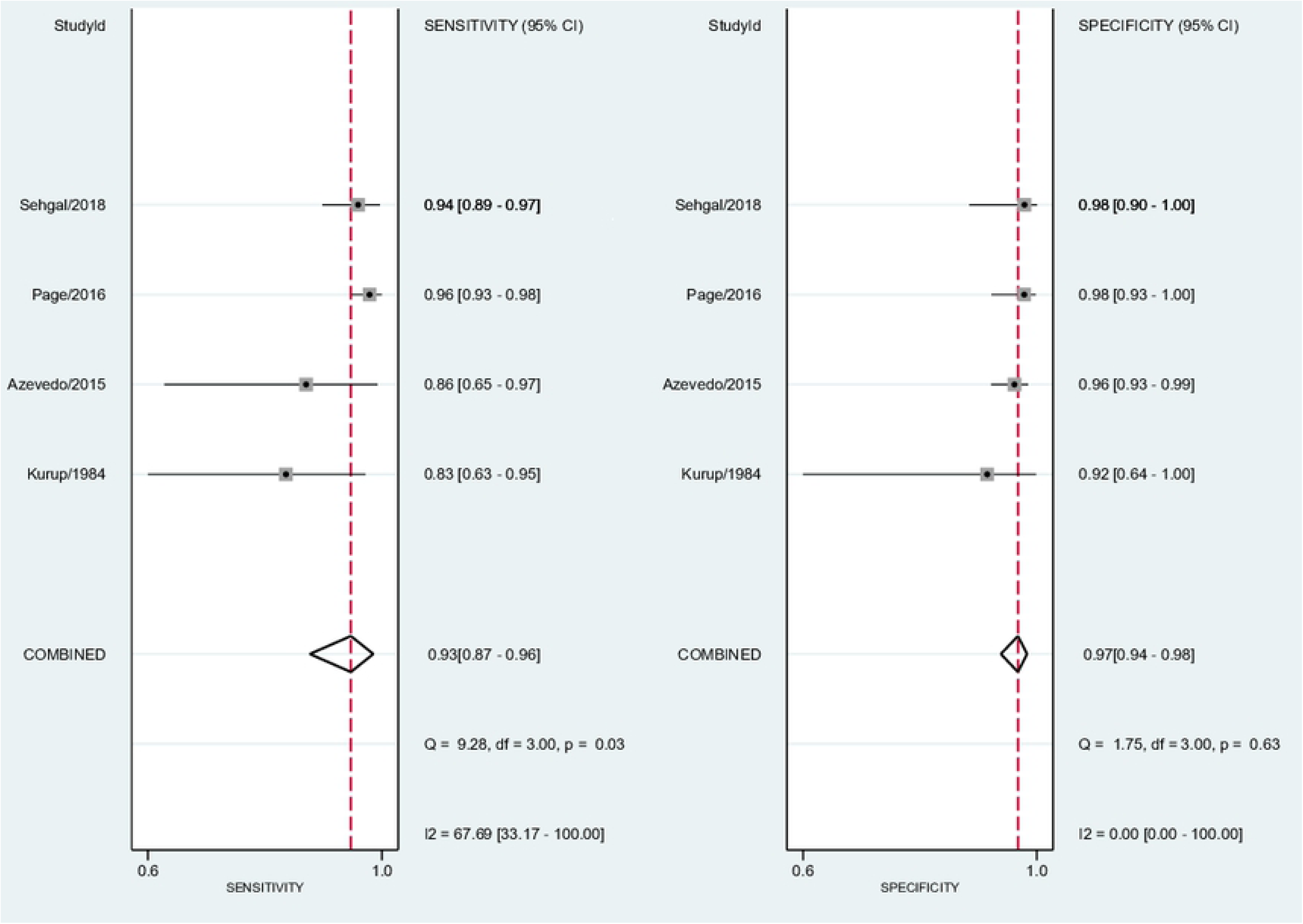
Forest plot for sensitivity, specificity and heterogeneity from four ELISA studies.

**Fig 5.**
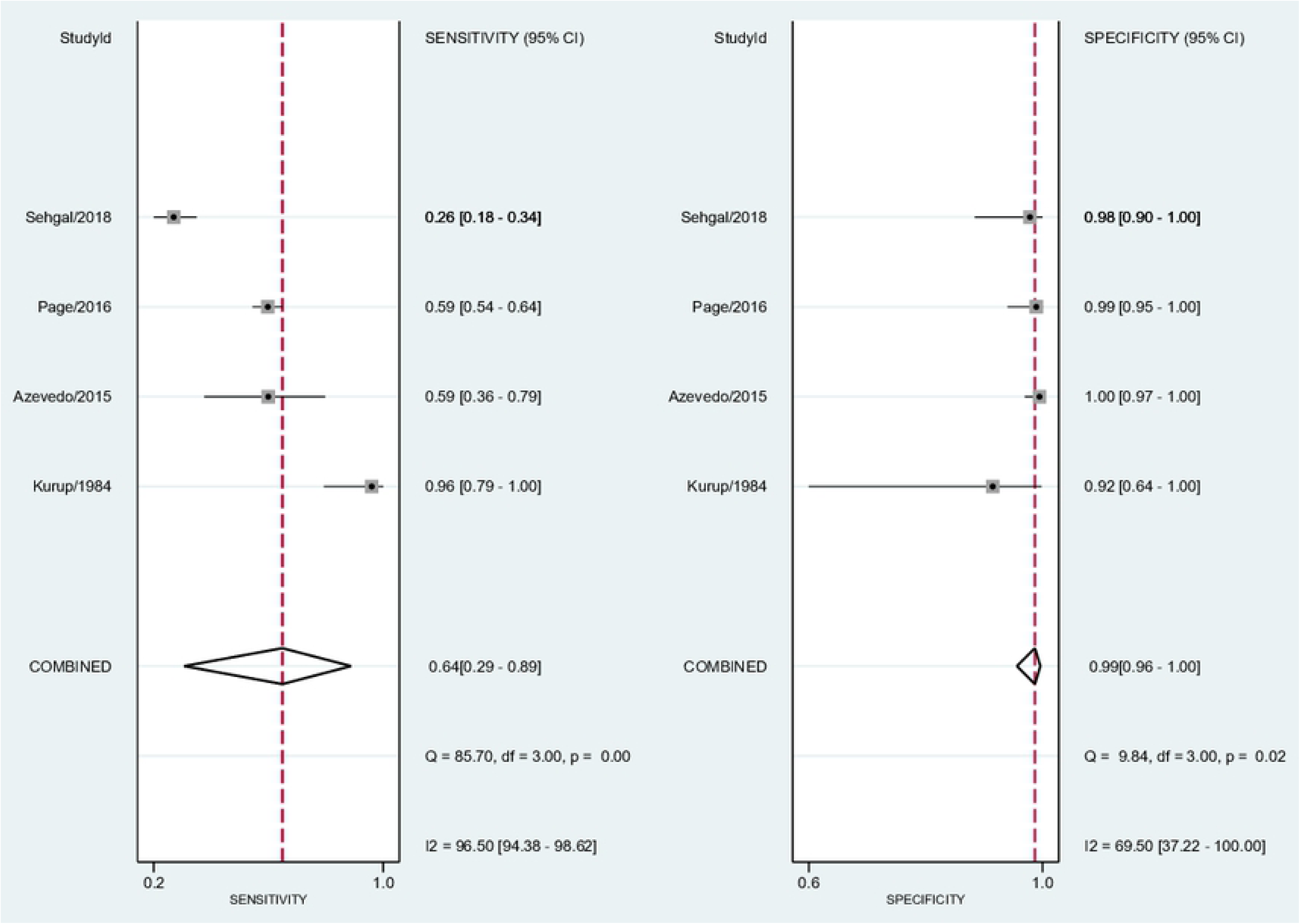
Forest plot for sensitivity, specificity and heterogeneity from four DID and/or CIE studies.

We also constructed the sROC curves and calculated the area under ROC (AUROC) for included studies (Fig 6). The overall diagnostic performance of the ELISA and the reference test (DID and/or CIE) were comparable (AUROC 0.99 [95% CI 0.97 to 0.99], and 0.99 [95% CI 0.97 to 0.99], respectively).

**Fig 6.**
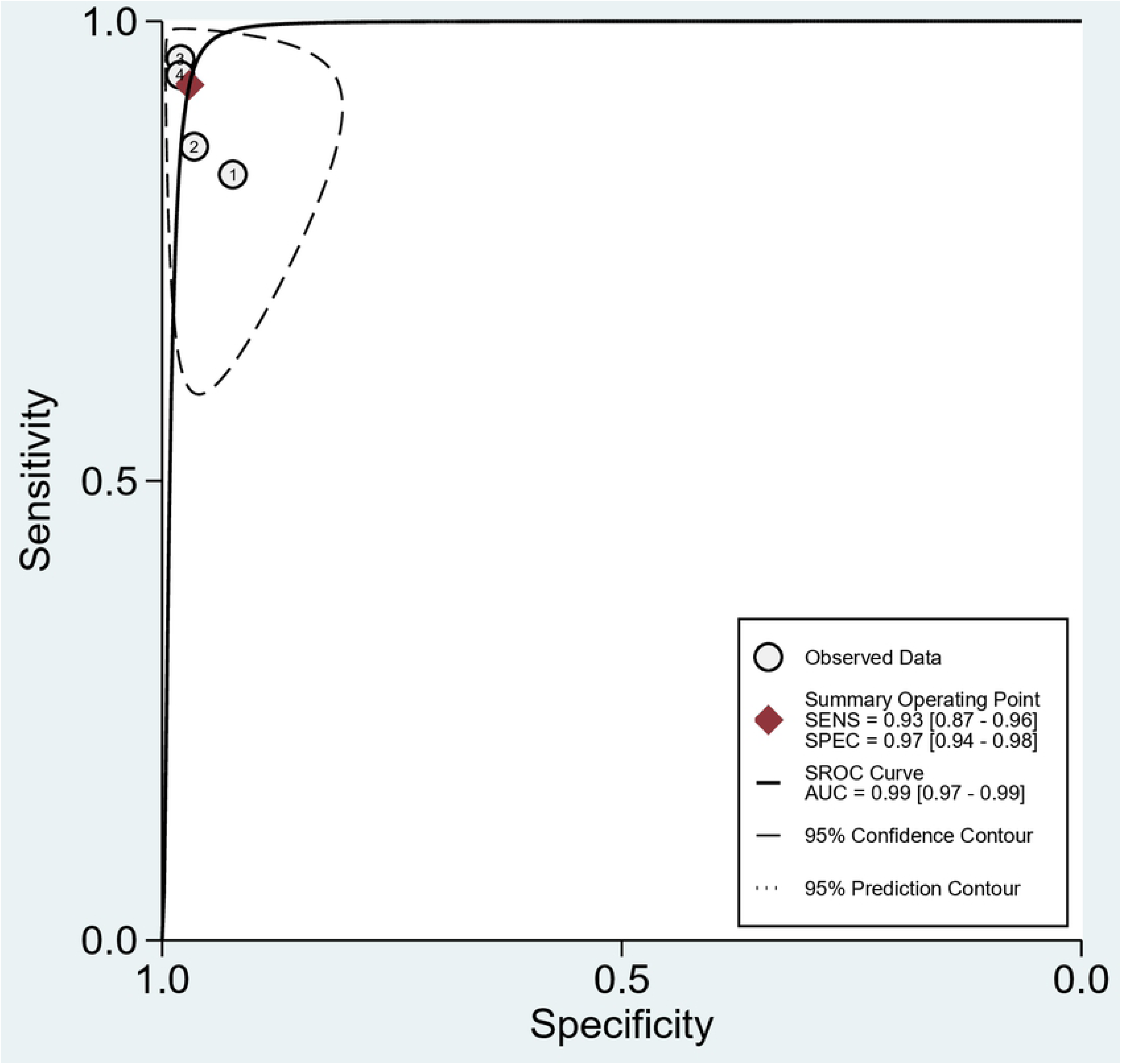
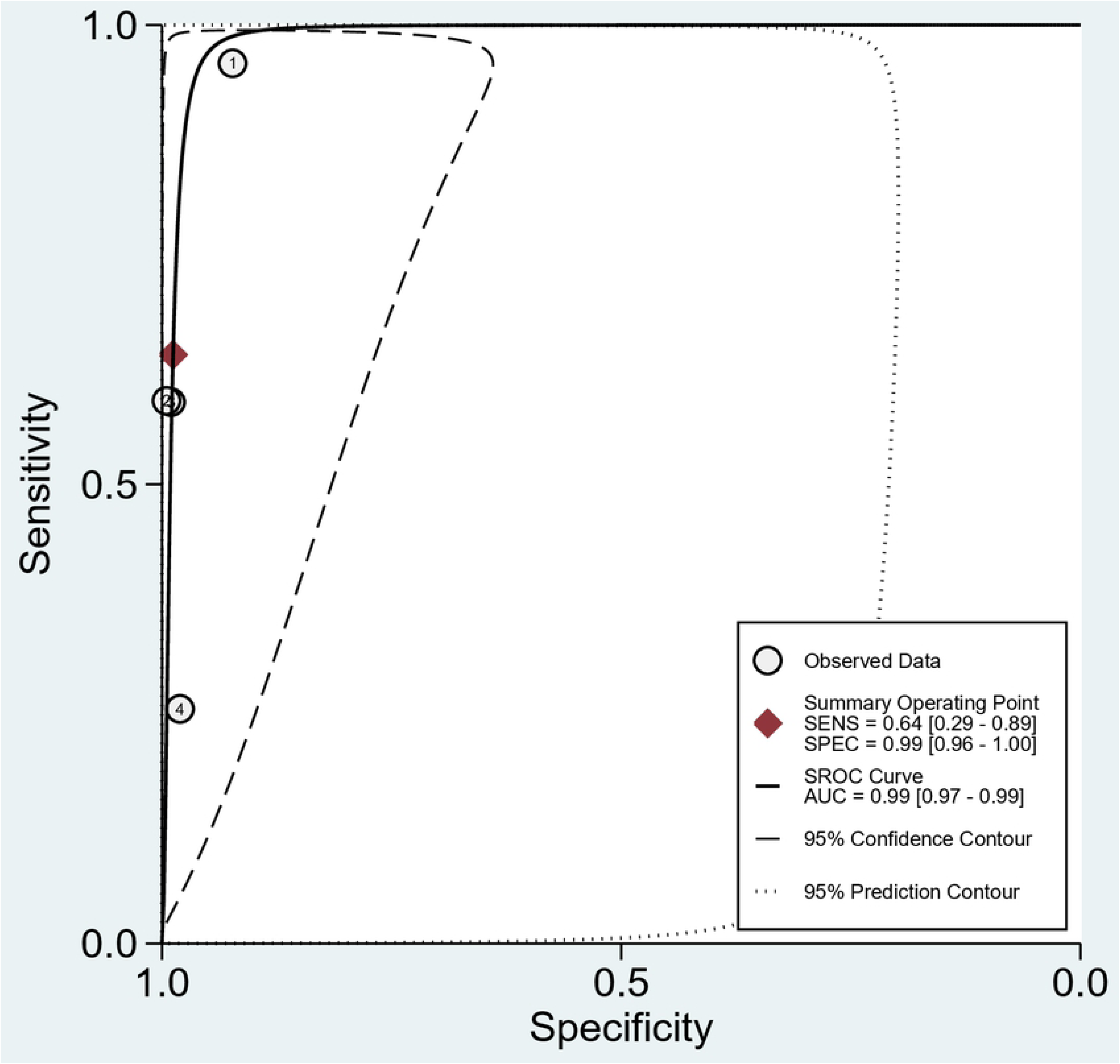
Summary ROC curves from the four included studies. A. AUROC for the ELISA test; B. AUROC for the reference test (DID and/or CIE).

### Investigations of heterogeneity

When we evaluated the four studies [22, 23, 24, 27], we found a heterogeneity (I2) of 67.69 (95% CI 33.17 to 100.00) in the ELISA sensitivity pool, considered as moderate heterogeneity, and 96.50 (95% CI 94.38 to 98.62) in the DID and/or CIE sensitivity pool, considered to be highly heterogeneous. We investigated the subgroup analyses, evaluating only the two most recent studies using commercial ELISA tests [24, 27] and the heterogeneity (I2) was 0% for sensitivity and specificity. When we studied the reference tests, the heterogeneity (I2) was 97.8% for sensibility and 0% for specificity.

The pooled sensitivity and specificity for the ELISA test, based on two data studies [24, 27], were 0.95 (95% CI 0.93 to 0.97) and 0.98 (95% CI 0.95 to 1.00), respectively. Pooled LR+ and LR- were 54.92 (95% CI 16.08 to 187.64) and 0.05 (95% CI 0.03 to 0.07), respectively. Pooled DOR were 1231.40 (95% CI 326.00 to 4651.70). The pooled sensitivity and specificity for the reference test (DID and/or CIE), based on two data studies [24, 27], were 0.49 (95% CI 0.45 to 0.54) and 0.99 (95% CI 0.96 to 1.00), respectively. Pooled LR+ and LR- were 55.39 (95% CI 7.82 to 392.60) and 0.56 (95% CI 0.29 to 1.06), respectively. Pooled DOR were 100.07 (95% CI 11.84 to 845.84). These results are presented in Figs 7 and 8.

**Fig 7.**
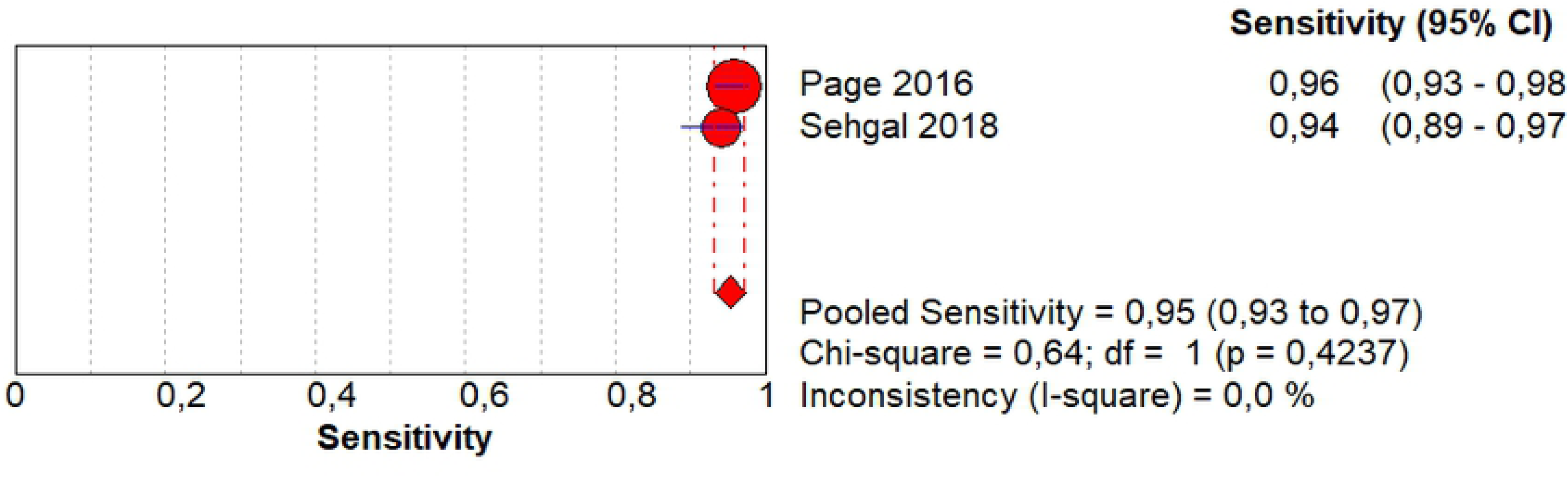
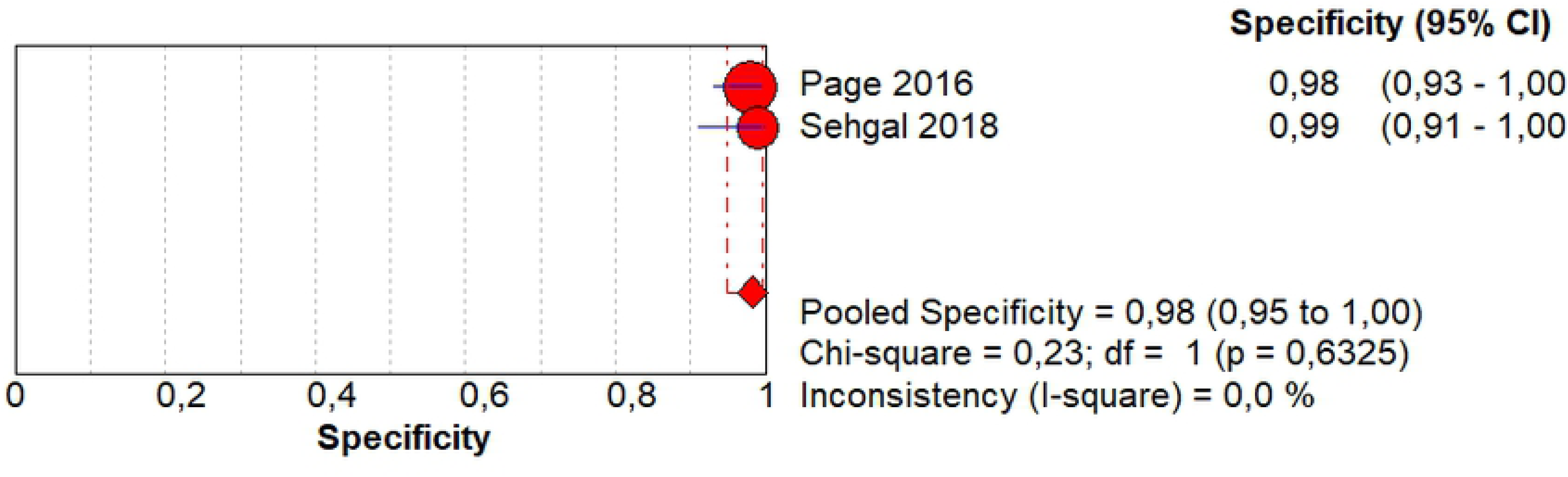
Forest plot of sensitivity (A), specificity(B) and heterogeneity from the ELISA test for the subgroup analyses (two studies).

**Fig 8.**
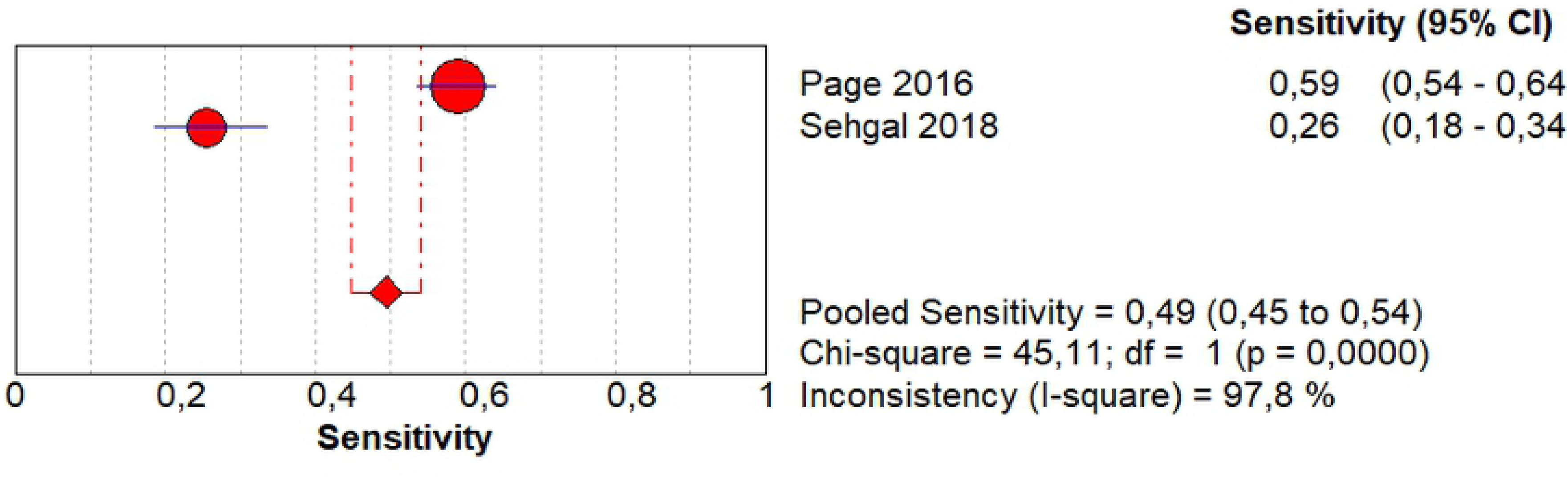
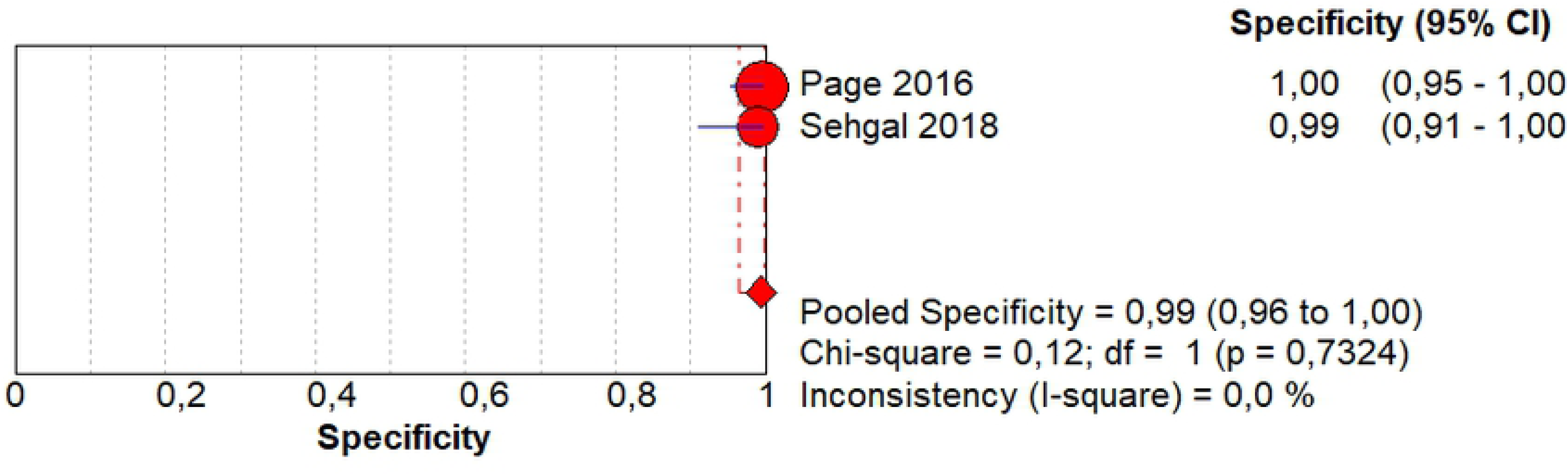
Forest plot of sensitivity (A), specificity (B) and heterogeneity from the DID and/or CIE test for the subgroup analyses (two studies).

Studies using in-house ELISA tests show large methodological differences in their performance. In the DID and/or CIE tests, high heterogeneity was maintained for the sensitivity in both studies [24, 27], considering that the precipitation tests are all in-house and also present large methodological differences in the studies included in this review.

## Discussion

This is the first systematic review comparing the ELISA test with the precipitin tests (DID and/or CIE) for the diagnosis of chronic pulmonary aspergillosis. Although current studies suggest ELISA as a better performance test for CPA diagnosis, precipitation tests are still considered in many countries as the reference test, especially in Brazil, where this review was carried out.

Thirteen articles that met the criteria for the research question were included, and all studies were considered as having an uncertain or high risk of bias in some domains in the quality risk assessment.

Important methodological differences were verified, mainly related to the in-house ELISA tests. More recent studies with commercial ELISA tests were included in the review, but also with differences described. We also observed this phenomenon in the DID and/or CIE tests, as these are all still in-house.

Mainly in former studies, we observed that the population selection was based on stored samples from patients already diagnosed with CPA and submitted to the tests described in the review. In addition, the lack of a checklist in the studies’ description was very evident, where many items in QUADAS-2 were not reported clearly, interfering with the quality of the evaluation. As an example, we noted that, in one study, although we were skilled in extracting the data for constructing the 2 x 2 table, the discussion and conclusion of the study had an error in printing and they were not compatible with the objective, methods and results of the article [22].

In the ELISA evaluation in individual studies included in the meta-analysis, the best performances based on the Youden’s test were from the commercial tests [24, 27], with ImmunoCAP and Immulite tests, ranging from 0.94 to 0.96.

When we evaluated Youden’s J statistic for the precipitation test (DID or CIE), in the studies included in the meta-analysis, only one study presented a performance of 0.96 [22] and the other studies [23, 24, 27] ranged between 0.26 and 0.59.

In a review article [35] it was reported that precipitin tests do not detect all CPA cases, but are correlated with disease activity and may become negative, so they can function as a follow-up tool along with imaging and inflammatory markers.

The ELISA test seems to be a promising test, and even with important methodological differences, it was useful to evaluate the use of diagnostic data for chronic lung aspergillosis in each study where it was possible to obtain data for the calculation of sensitivity and specificity. Two more recent studies were highlighted in this review [24, 27], with sensitivities presenting lower confidence intervals for the ELISA test, and when compared to the confidence intervals from the reference tests (DID and/or CIE), they showed a better performance. Besides that, the pooled LR+/LR- from the ELISA test presented as conclusive evidence and this was not observed in the reference test results.

Several studies have recently been published with serological data using only commercial ELISA tests for CPA diagnosis in an area with a high prevalence of tuberculosis [1, 13, 36].

The limitations of this study rely in the primary studies. There were problems regarding individual reporting for the primary studies, thus we could not do a 2×2 table; in some cases the lack of appropriate reporting made us judge the study as having an unclear [22, 28, 30, 33, 34] or high risk of bias [31].

The availability of commercial tests demonstrated in recent studies [24, 27] may facilitate the incorporation of the ELISA test into our clinical practice, allowing standardized use for the diagnosis of chronic pulmonary aspergillosis and replacing the reference test that still depends on its in-house performance.

Because the global burden of CPA is substantial, mainly as a sequel to pulmonary tuberculosis (PTB) [37] and especially in countries such as Brazil, which is on a list of 30 countries representing over 80% of tuberculosis cases worldwide in 2015 [38], there is still a need for well-designed studies so that the degree of evidence is obtained and demonstrated for the use of the ELISA test in comparison to the precipitation tests.

In conclusion our meta-analysis suggests that the enzyme-linked immunosorbent assay (ELISA) presented a better accuracy than the precipitation tests (DID and/or CIE) for CPA diagnosis, and that it can be considered the test of choice in clinical practice.

## Supporting information captions

**S1 Table.**
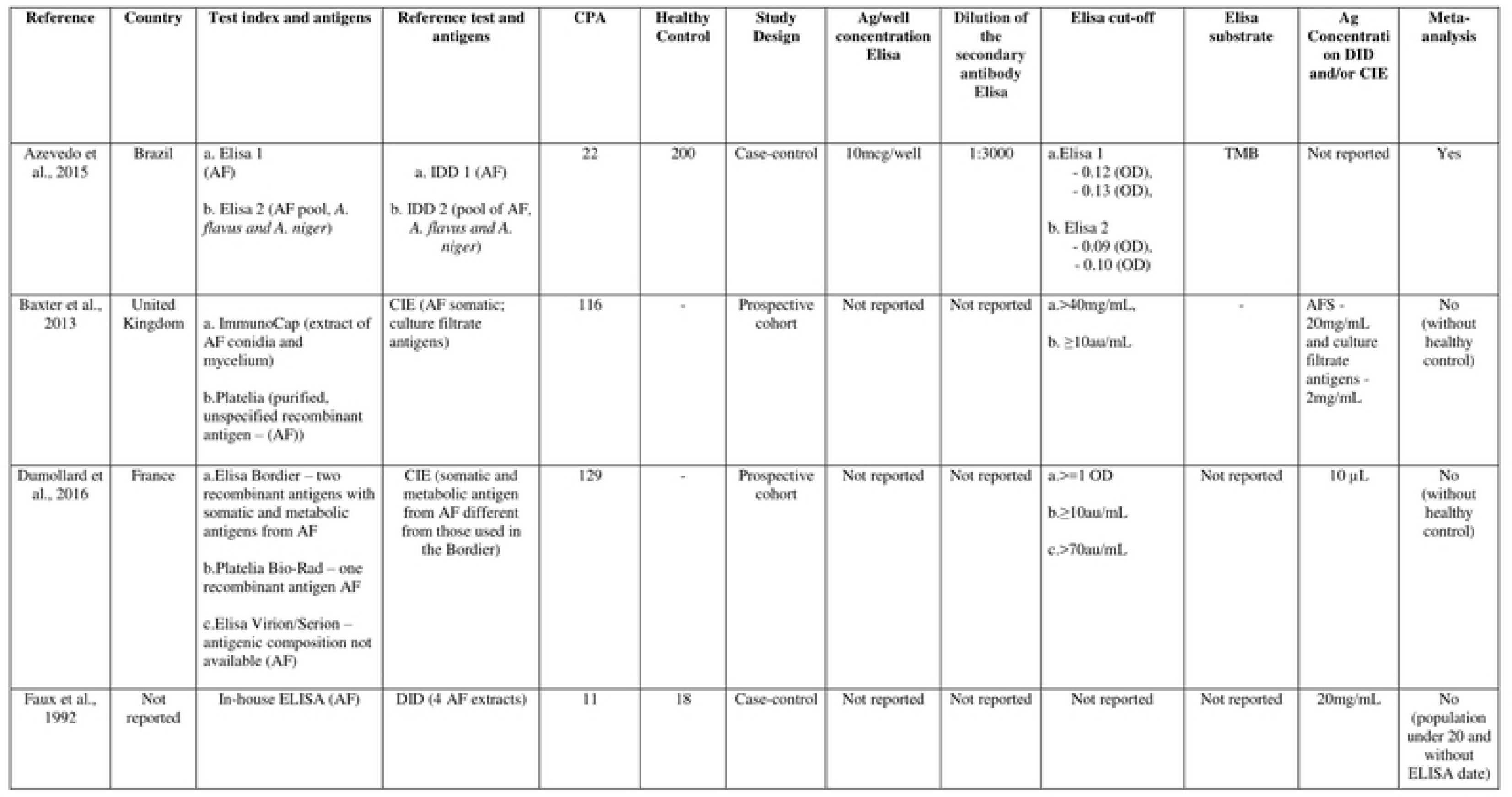
Characterization of the studies included in this systematic review and meta-analysis. ELISA: Enzyme-Linked Immunosorbent Assay; AF: *Aspergillus fumigatus*; Ag: antigen; DID: Double Immunodiffusion; CPA: chronic pulmonary aspergillosis patients; OD: optical density; CIE: counterimmunoelectrophoresis; TMB: 3,3′,5,5′-Tetramethylbenzidine; pNPP: Alkaline Phosphatase Yellow; OPD: o-Phenylenediamine; RNU: 18-kDa ribonuclease; DPPV: 88-kDa dipeptidylpeptidase; CAT: 360-kDa catalase

